# The Arctic copepod *Calanus hyperboreus* is more tolerant to marine heatwaves than temperate copepods in the Oslofjord

**DOI:** 10.1101/2025.09.21.677595

**Authors:** Mathieu Lutier, Andrea Emilie Thorstensen Skari, Nele Thomsen, Helena C. Reinardy, Khuong V. Dinh

**Author notes:** Correspondence and requests for materials should be addressed to ML.

## Abstract

*Calanus hyperboreus* plays a key role in the functioning of Arctic ecosystems. It is considered highly vulnerable to ocean warming (OW) and marine heatwaves (MHW), which would reduce its range, expected to shift northward. Yet, *C. hyperboreus* is reported as far south as the Skagerrak, where it is considered non-native and transported by ocean currents. We argue that this may be an isolated population adapted to warmer temperatures. To test this hypothesis, we exposed *C. hyperboreus* from the Oslofjord to temperatures from 0 to 24 °C, for 5 days. We recorded survival to identify the upper threshold of thermal tolerance and DNA damage to detect sublethal effects. The thermal response of *C. hyperboreus* was compared with that of the dominant copepod species in the Oslofjord, *Calanus finmarchicus* and *Metridia longa*. We found that the survival of *C. hyperboreus* did not decrease before reaching 16-20 °C which was much higher than 13-16 °C and 4-8 °C for *C. finmarchicus* and *M. longa*, respectively. *C. hyperboreus* showed the least DNA damage, highlighting the adaptation of its physiology to the Oslofjord. Our results suggest the existence of local adaptations to warming in *C. hyperboreus* that could determine its fate under climate change.

## Introduction

Climate change will cause ocean temperatures to increase by 1–5 °C by the end of the 21st century, a phenomenon called ocean warming (OW) [1,2]. The Arctic Ocean is the most vulnerable, warming 4–7 times faster than the global ocean average [3]. Marine heatwaves (MHW) are periods when surface waters are much warmer than the normal seasonal average, up to + 5 °C, which can persist for months [5]. Climate change makes MHW exponentially more frequent [4]. Temperature is one of the main parameters controlling the biology of ectothermic organisms [5,6]. Each organism has a temperature optimum and an upper tolerance threshold beyond which its homeostasis and survival are compromised [7,8]. Thus, OW and MHW are restricting the habitats and shifting the distribution of many species [9– 11]. This causes a poleward shift of species distributions and reduces the range of polar species, threatening some of them with extinction [12,13].

A striking example is the Arctic copepod *Calanus hyperboreus* (Krøyer, 1838), which could experience a northward shift of its distribution by up to 70 km per decade [14]. *C. hyperboreus* presents unique features in copepods being one of the largest, up to 7 mm prosome length, and the richest in lipids, up to 60% of its dry body weight (DW) [15–17]. It is the preferential prey of higher trophic levels, particularly Arctic fish and seabirds, in regions where it dominates zooplankton [15]. After storing lipid reserves over spring/summer, *C. hyperboreus* migrates to the deep sea where it overwinters at 600-3000 m depth [15,17]. By natural mortality and respiration, this sequesters a huge amount of carbon in the deep sea [18]. Therefore, changes in the distribution of *C. hyperboreus* would have large consequences for ecosystems.

*C. hyperboreus* shows highest abundances in waters with temperature ranges between 0 and 5°C and is rare above 7°C [19–21]. The upper thermal tolerance threshold for its survival has been determined at 10-15 °C [22,23]. Thus, the heart of the species’ distribution range is in Arctic regions and mainly in the Greenland Sea, where the highest abundances are recorded (Figure 1) [15]. Yet, *C. hyperboreus* is observed as far south as the Skagerrak at the mouth of the Baltic Sea (Figure 1) [24–29]. In these areas, the upper tolerance limit of 10-15°C is commonly exceeded [30] which is amplified by MHW [31]. *C. hyperboreus* is considered a non-native species to the North Sea, where it is thought to be transported from the north by ocean currents [24,27,32]. Overall, the species abundance is negligible in the Norwegian and North Seas (Figure 1). Yet, data from recent cruises found high abundances, up to 3.2 g DW m^-2^, of *C. hyperboreus* in the deepest part of the Skagerrak (∼600 m) [28,29]. Such abundance is similar to those reported in many Arctic regions such as the Makarov, Canada, Nansen, and Amundsen Basins [33–36]. This highlights a clear disconnection in *C. hyperboreus* distribution with negligible abundances between the Iceland Sea and Skagerrak. The isolation of the Skagerrak *C. hyperboreus* population is also supported by the smaller size of its genome in comparison with Arctic conspecifics [25]. Therefore, we hypothesize that *C. hyperboreus* from Skagerrak is an isolated population adapted to warmer water temperatures.

**Figure 1.**
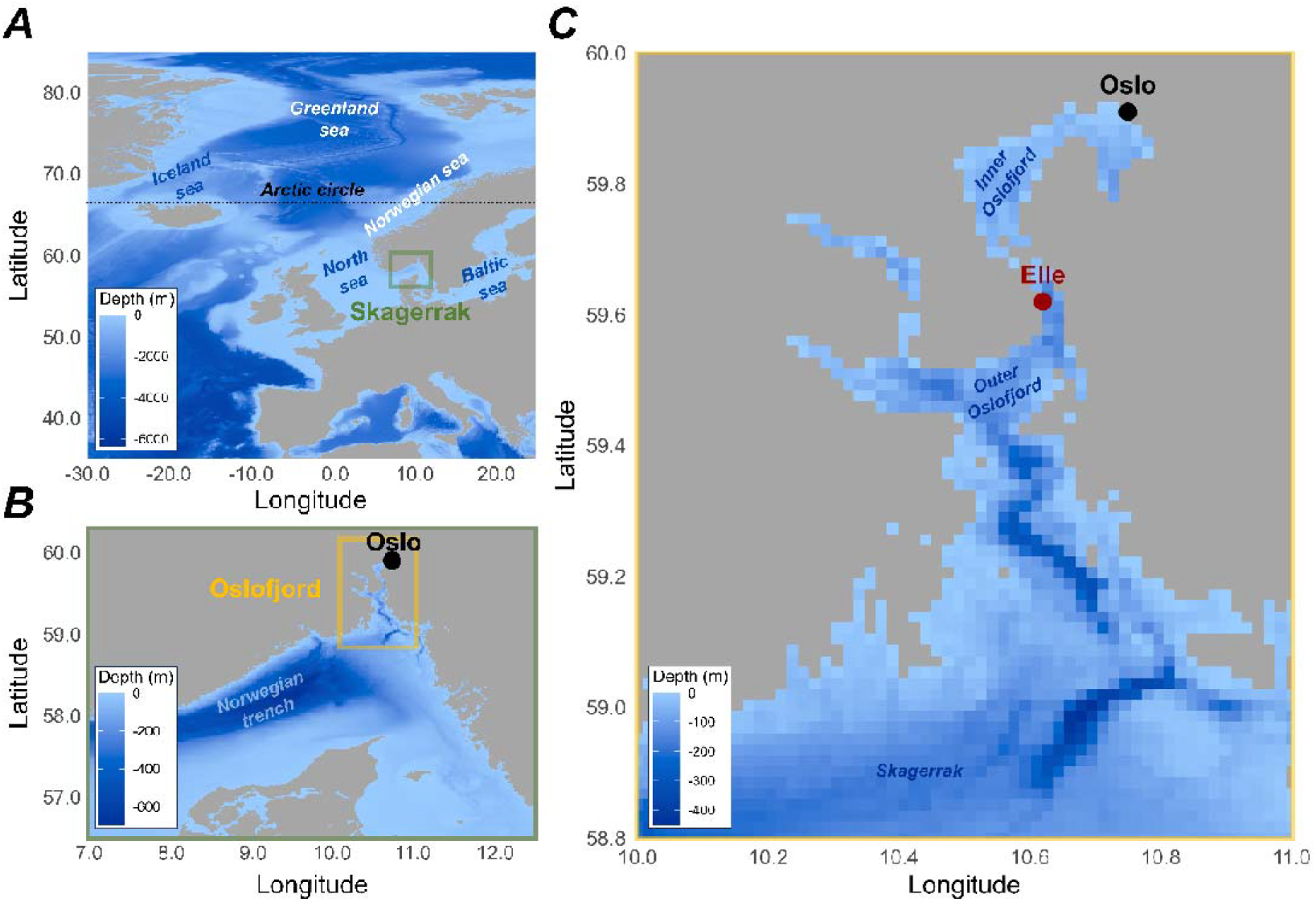
Geography of the Oslofjord, the study location. A) The position of the Skagerrak in the North Sea, a subregion of the Atlantic Ocean, is described (green square). B) The position of the Oslofjord in the Skagerrak is described (yellow square). C) The geography of the Oslofjord is described, as well as the location of Elle, the sampling station (red dot). The positions of major seas, the Arctic Circle (dotted line), and points of interest cited in the article are indicated with text, to be used as spatial references.

To test this hypothesis, we exposed *C. hyperboreus* collected in the Oslofjord (North Skagerrak) to temperatures from 0 to 24 °C. Survival was recorded for 5 days to determine the upper thermal tolerance threshold. To detect sublethal effects, we also measured the amount of DNA damage that can reflect oxidative stress caused by acute heat shock [37]. For reference, we compare the thermal response of *C. hyperboreus* with that of the dominant copepod species in the Oslofjord at the time of sampling, *Calanus finmarchicus* (Gunnerus, 1770) and *Metridia longa* (Lubbock, 1854). *C. finmarchicus* and *M. longa* are commonly observed in the Oslofjord and we expect them to be more tolerant to warming. Determining the existence of a heat-tolerant population is crucial because local adaptations can provide a reservoir of resistant genotypes during rapid environmental changes [38,39].

## Material & methods

### Sampling

The Oslofjord copepod community was sampled on 5 June 2023, near Elle Station (59.62 °N, 10.62 °E) (Figure 1). The Oslofjord is the northern branch of the Skagerrak, the strait forming the border between the North Sea and the Baltic Sea (Figure 1). Sampling was carried out using a WP2 net (200 μm mesh) with three vertical hauls from the bottom (203 m) to the surface. The seawater temperature below the thermocline, where most calanoid copepods are distributed during the day [40,41], was recorded with a CTD and was approximately 8 °C. Samples were then transported to the laboratory at the University of Oslo, where copepod species were sorted at ∼8 °C, under a microscope, in a climate room. Species and life stages were identified based on morphological criteria and the number of segments on their urosome. Copepodite 5 (CV) and adult females (AF) were kept for the experiment at a ratio of 69% and 31%, respectively. The other two dominant species were *M. longa* (51% AF, 46% adult males, AM, and 3% CV) and *C. finmarchicus* (87% CV, 9% AF, 3% AM, 1% CIV). For each species, we used a mix of life stages (as presented above) that represented the population demography at the time of sampling. After sorting, copepods were maintained in opened bottles of filtered seawater at a density of 45 copepods/L in an incubator at 8 °C. The same batch of seawater was used to maintain copepods throughout the experiment. This seawater was collected on 5 June 2023 at Drøbak Aquarium (pumped at 40 m depth in the Oslofjord). The seawater was filtered at 1 μm, treated with a UV lamp, and stored at 4 °C. Air was bubbled into the seawater to ensure good oxygenation. Copepods were not fed during the experiments. All three species have large lipid reserves to survive the low-feeding season [18,42,43], and these are unlikely to have decreased during the short duration of the experiment. This was verified by lipid droplet measurements (see below).

### Experimental design

*C. hyperboreus, C. finmarchicus*, and *M. longa* were exposed to seven different temperatures: 0, 4, 8, 12, 16, 20, 24 °C. These conditions cover the range of temperatures recorded on the whole water column at the sampling station during monthly measurements in 2020 that ranged from 3-20 °C (Even Sletteng Garvang personal communication). The 24 °C condition was selected to test the influence of future OW and MHW. Each temperature condition was maintained in one of seven incubators in which copepod exposures were performed.

On 14 June 2023, individual copepods were placed in plastic flasks (VWR^®^ tissue culture flasks) filled with 65 mL of seawater. 35 *M. longa*, 15 *C. finmarchicus*, and 5 *C. hyperboreus* were then placed in each of the 7 incubators set at an initial temperature of 8 °C (Figure 2). The different sample sizes for the three species are due to the different abundances found in the field sample. For 4 days, the temperature was then increased (for conditions above 8 °C) or decreased sequentially (for conditions below 8 °C) by 4 °C d^-1^ so that all final temperature conditions were reached on the same day in the different incubators (Figure 2). The exposure lasted 5 days because it is the minimum duration of a MHW in the environment [44]. Survival was checked daily by gently tapping on the wall of the flask with a pin to stimulate the copepods, and immobile copepods were considered dead. Temperature was recorded in each incubator every 15 minutes using HOBO^©^ TidbiT v2 loggers (1 logger/incubator) placed in a 65 ml beaker, sealed with parafilm, and filled with seawater. The average temperature in each incubator during the exposure period was calculated and used for statistics. Photographs of copepods were taken daily, before removal, for dead copepods, and at the end of the experiment, before sampling, for survivors, using a stereomicroscope (Leica M 125 C, LAS X software). Prosome size, and lipid droplet and prosome surface areas were measured using Image J. Lipid reserve (%) was determined as lipid / prosome area.

**Figure 2.**
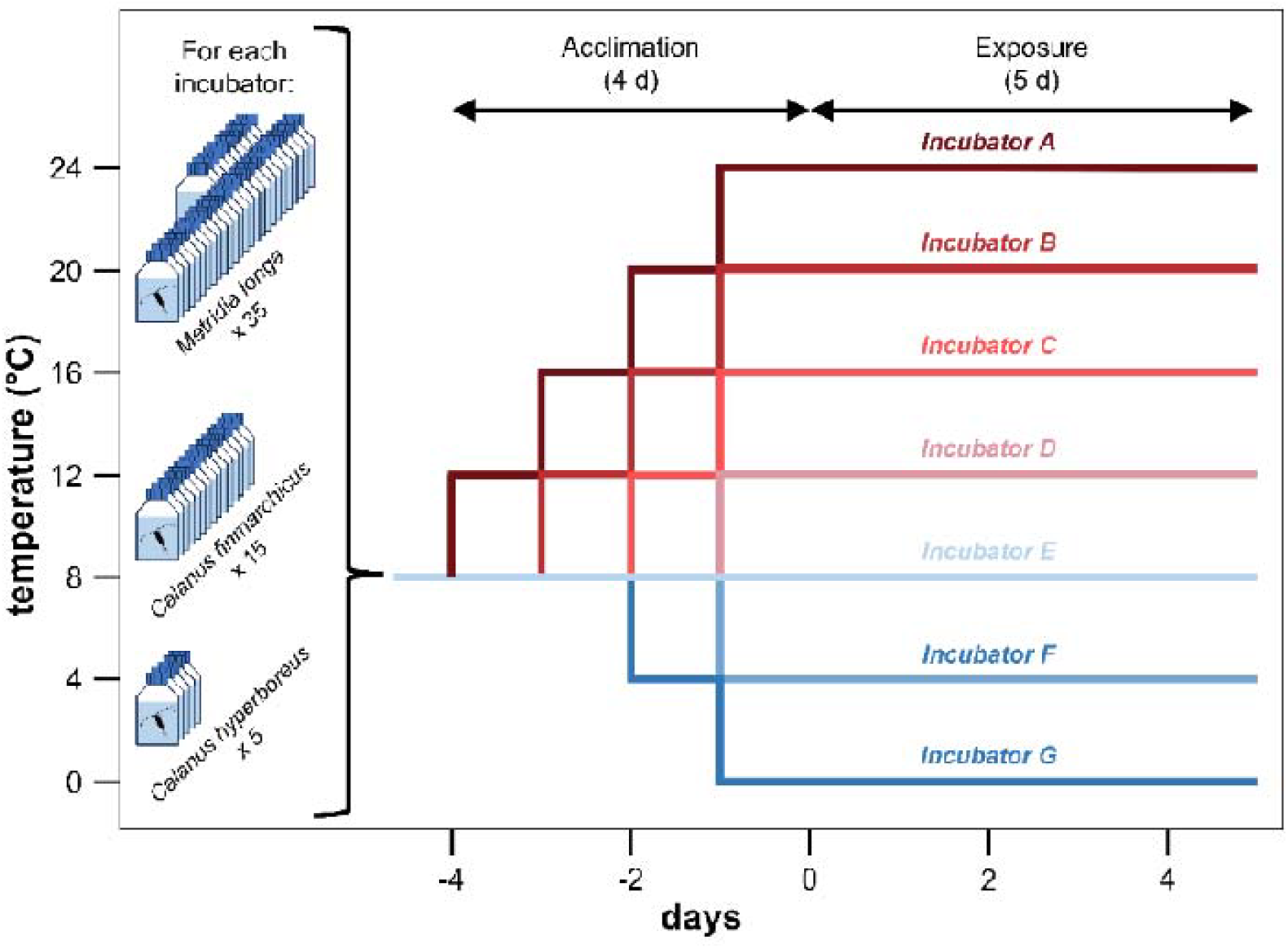
Experimental design used to determine the temperature tolerance of three species of copepods from the Oslofjord: *Metridia longa, Calanus finmarchicus* and *Calanus hyperboreus*. Variations in temperature over time are presented for the 7 incubators, each containing 35 *M. longa*, 15 *C. finmarchicus* and 5 *C. hyperboreus*. The 4 first days corresponded to the gradual changes of temperature in the incubators. The exposure then lasted 5 d for all the temperature conditions.

### DNA damage analyses

The Fast MicroMethod assay (FMM) was used for determining the amount of DNA damage by assessing the unwinding kinetics of double-stranded DNA in alkaline solution with fluorescence measurements [45,46]. For each species, at the end of the experiment, 5 ± 1 copepods (depending on the final survival, Supplementary Table 1) from each temperature condition were individually snap**-**frozen in liquid nitrogen and stored at −80 °C. Our protocol is adapted from Halsband *et al*. [47]. Individual copepods were gently homogenised with a hand pestle in 260 μL of 20 mM EDTA + 10% DMSO. 20 μL of homogenate, or 20 mM EDTA + 10% DMSO for the blank, was loaded, with 5 analytic replicates per sample, into a microplate (96-well black-walled, Greiner Bio-One Ltd). Samples were lysed on ice for 40 min in the dark after adding 20 μL of lysis buffer to each well (20µl 9M urea, 0.1% SDS, 0.2M EDTA, 2% Quant-iT™ PicoGreen®). After lysis, unwinding was initiated by adding 200 μL of unwinding solution (mixture of 20 mM EDTA and 2 M NaOH until pH is adjusted to 13.0) to each well. Fluorescence was then recorded immediately using a SynergyMx^®^ microplate reader (BioTek^©^) and every minute for 30 min (excitation 485nm, emission 520nm).

We estimated the amount of DNA damage for each sample as the Slope of the Linear Regression (SLR, RFU min^-1^) of fluorescence with time [48,49]. This is different from the strand scission factor (SSF) which is commonly used to estimate the amount of DNA damage, but was less suitable for our experimental design (see more in Supplementary Note 1). An increase in SLR shows an increase in DNA damage. The linear regression of fluorescence over unwinding time was fitted between 0 and 20 minutes, representing the linear part of the DNA unwinding. We only retained significant linear regressions (*p < 0*.*05*), which respected the assumptions of normality and homoscedasticity of the residuals. The 3 best analytic replicates were kept for each sample using z-scores, i.e. the three replicates that diverged the less from the mean SLR of the 5 replicate wells [50]. The mean SLR was then calculated from these 3 replicates for each sample, and kept for statistics.

### Statistics

Analyses were performed using the R software v4.3.1 and the statistical significance threshold was 0.05. The effects on survival of temperature, species, prosome length, lipid reserves, and the interaction of all these factors, were tested using Cox proportional-hazards model using the *coxme* function of the *coxme* package [51]. We did not include DNA damage as a factor influencing survival because it is not measurable in dead copepods. The different factors were tested and the model with the lowest Akaike information criterion (AIC) and Bayesian Information Criterion (BIC) values was retained. Therefore, only temperature and species were kept while size and lipid reserves did not affect survival. The assumption of proportional hazards of the Cox model was graphically checked by plotting the Schoenfeld residuals against time using the *ggcoxzph* function of the *survminer* package [52]. For significant factors, i.e. temperature and/or species, *post-hoc* pairwise comparisons were performed using the *pairwise_survdiff* function. Sample sizes are unbalanced for the three species (n = 245 for *M. longa*, n = 105 for *C. finmarchicus* and n = 35 for *C. hyperboreus*). Cox models are valid for an unbalanced design, but different samples sizes affect the statistical power, i.e. the ability to detect an effect of the factors [53–55]. To test differences in survival response of species at different exposure temperatures, a first general model was built for the three species, with an unbalanced design. Then individual models were tested for each species separately, with a balanced design. Kaplan-Meier survival curves were used to visualize survival as a function of time. Type II ANOVA, for unbalanced designs (n = 28 for *M. longa*, n = 29 for *C. finmarchicus*, n = 24 for *C. hyperboreus*, Supplementary Table 1), was used to test the effects of temperature, species, and their interactions on DNA damage (SLR). Size and lipid reserves did not affect DNA damage. The homogeneity of variances and normality of residuals were checked graphically. The differences between groups for significant factors were tested using Tukey HSD tests.

## Results

Temperatures in the incubators were stable during the 5 days of exposure and were 0.0 ± 0.4 °C, 3.9 ± 0.5 °C, 7.9 ± 0.2 °C, 13.3 ± 0.2 °C, 16.0 ± 0.2 °C, 19.9 ± 0.3 °C, and 23.8 ± 0.2 °C (mean ± standard deviation, N = 670). The survival curves were significantly different for species (*p < 0*.*001*) and temperatures (*p < 0*.*001*) (Cox proportional-hazards model, Supplementary Table 2). Survival of *M. longa* was generally lower than those of *C. finmarchicus* and *C. hyperboreus*, which were not significantly different (*post-hoc* tests, Figure 3). The interaction of the two factors was significant (*p < 0*.*001*), indicating that the response to temperature was different for the three species. Individual models for each species confirm that temperature significantly decreased the survival of *M. longa* (*p < 0*.*001*, Figure 3A), *C. finmarchicus* (*p < 0*.*001*, Figure 3B) and *C. hyperboreus* (*p < 0*.*001*, Figure 3C).

**Figure 3.**
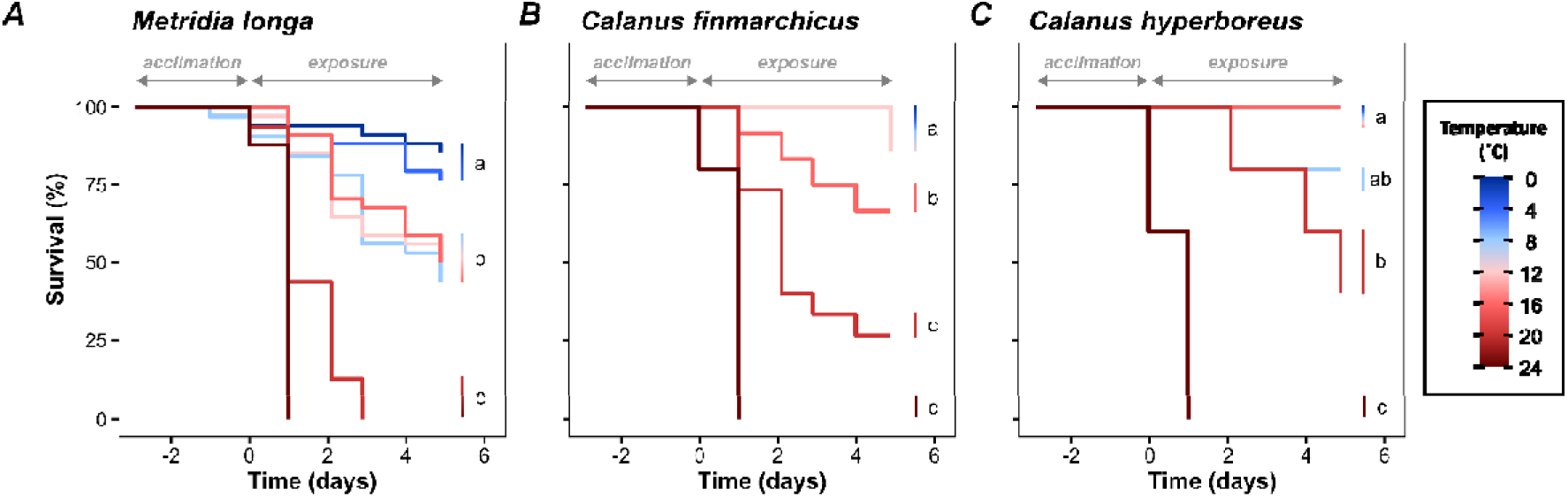
Survival of three copepod species from the Oslofjord as a function of time and temperature. Kaplan-Meier survival curves are shown for A) *Metridia longa*, B) *Calanus finmarchicus*, and C) *Calanus hyperboreus*. The temperature during the exposure period is represented by the colour gradient. Survival curves that differ significantly for different temperatures are indicated by different letters. Some survival curves overlap, this is the case for B) temperatures 0.0, 3.9, 7.9 and 13.3 °C and C) 0.0, 3.9, 7.9, 13.3 and 16.0 °C. Negative values for time on the x-axis indicate the days before the final temperature condition is reached, i.e. the “acclimatisation” period. The exposure period is indicated by positive values.

*M. longa* showed no difference in survival between 0.0 and 3.9 °C with a final survival rate of 81 ± 4 % (post-hoc test, Figure 3A). Survival then significantly decreased at 7.9-16.0 °C with a final survival rate of 48 ± 3 %. Finally, there was no survival at 19.9 and 23.8 °C after 1 and 2.9 days of exposure respectively. *C. finmarchicus* showed no difference in survival between 0.0 and 13.3 °C with a final survival rate of 95 ± 6 % (post-hoc test, Figure 3B). Survival significantly decreased at 16.0 °C and 19.9 °C with final survival rates of 67 % and 27% respectively. No survival was recorded at 23.8 °C after 1 day of exposure. *C. hyperboreus* overall showed no difference in survival between 0.0 and 16.0 °C with a final survival rate of 96 ± 8 % (post-hoc test, Figure 3C). Survival then significantly decreased at 19.9 °C with a final survival rate of 40 %. No survival was recorded at 23.8 °C after 1 day of exposure.

Overall, regardless of temperature, DNA damage differed significantly for the three species (ANOVA type II, p < 0.001). DNA damage was higher in *M. longa* (-230 ± 164) than in *C. finmarchicus* (-426 ± 157) and lowest in *C. hyperboreus* (-556 ± 113) (Tukey’s HSD test, Figure 4). The results of type II ANOVA indicate that temperature (*p = 0*.*038*) significantly increased DNA damage for all three species (Supplementary Table 3). There is no interaction between temperature and species, indicating that the response of DNA damage to temperature was similar between species.

**Figure 4.**
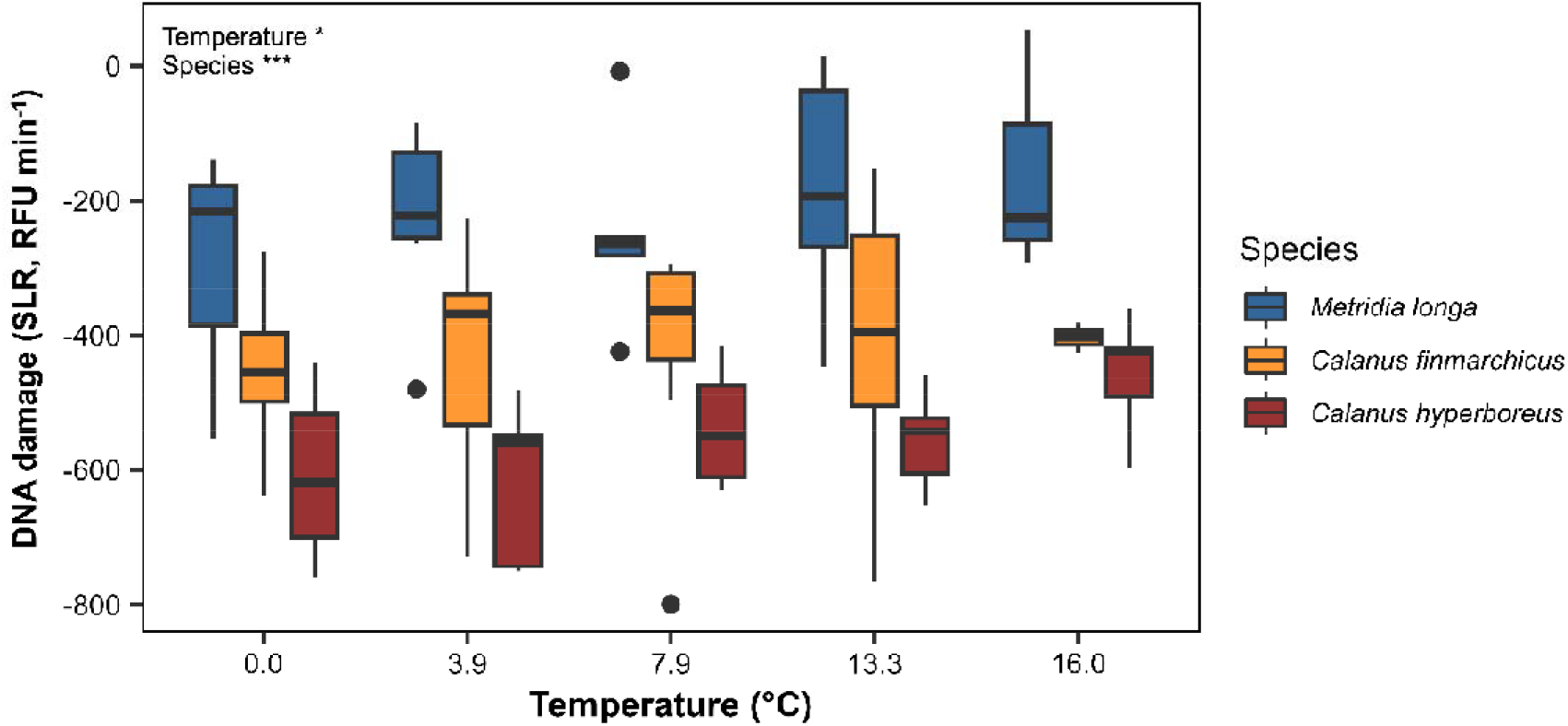
DNA damage in copepods from the Oslofjord that survived 5 days of exposure to different temperatures. Data are presented as box and whisker plots (averages and standard deviations). The species are *Metridia longa* (blue), *Calanus finmarchicus* (orange), *Calanus hyperboreus* (red). The results of type II ANOVA testing the effects of temperature, species and the interaction of the two factors on DNA damage are shown in the upper left corner.

## Discussion

We experimentally study for the first time the ecophysiology of *C. hyperboreus* sampled in the Skagerrak, ∼ 1500 km south of its Arctic range, where it is considered a non-native species, transported by ocean currents, with low resistance to warm temperatures. Unexpectedly, we find that *C. hyperboreus* is more resistant to warming than *M. longa* and *C. finmarchicus* which are commonly found in the Oslofjord. This suggests the existence of a local *C. hyperboreus* population in Skagerrak, adapted to withstand warm temperatures. This should be considered when studying OW and MHW impacts on the species.

In the Oslofjord, the survival of *M. longa* is most vulnerable to OW and MHW, much more than that of *C. finmarchicus*. We show that survival of *M. longa* decreases above 4-8 °C with no survival above 16-20 °C, while survival of *C. finmarchicus* decreases above 13-16 °C with no survival above 20-24 °C. *M. longa* is abundant in cold Arctic and boreal waters [56,57] and do not survive above 14-18 °C [22], with the Oslofjord being the southern limit of its distribution [58]. *C. finmarchicus* is a boreal species abundant between 0 and 10 °C [20,23], and do not survive above 20 °C [22]. If *C. hyperboreus* is a non-native species from the Skagerrak transported there by ocean currents, we expect it to be more sensitive to warming.

Contrary to expectations, we found that, in the Oslofjord, *C. hyperboreus* is more resistant to warming than *M. longa* and *C. finmarchicus*. Indeed, *C. hyperboreus* survival decreases above 16-20 °C with no survival above 20-24 °C. This is far greater than what is reported for Subarctic/Arctic *C. hyperboreus* that do not survive above 10-15 °C [22,23]. However, comparisons with literature should be made with caution since we use a different protocol. The strength of our study is comparing the tolerance of *C. hyperboreus* with the other species occurring in Oslofjord that should be adapted to warmer conditions. Because of different abundances in the field, we had 7 and 3 times fewer *C. hyperboreus* per condition than *M. longa* and *C. finmarchicus*, respectively. Therefore, the probability of having a high proportion of individuals surviving after 5 days was much lower for *C. hyperboreus*, and yet this occurred between 0 and 16 °C, suggesting our results are conservative. There was a difference in the proportions of life stages between the three copepod species used in the experiment. In some copepod species, older individuals survive stress better than younger individuals [59–61]. In our study, there were more adults in *M. longa* (51%) than in *C. hyperboreus* (31%) and significantly more than in *C. finmarchicus* (9%). Therefore, *C. hyperboreus* may have faced a disadvantage in terms of survival. Yet, it survived MHW much better than *M. longa*, confirming the robustness of our observations.

We find that *C. hyperboreus* exhibits less DNA damage than *C. finmarchicus*, with both species exhibiting less DNA damage than *M. longa*. DNA damage occurs continuously in organisms due to exogenous (e.g. UV, environmental stressors) or endogenous (production of reactive oxygen species, ROS, by metabolism) factors [62]. At the same time, organisms continually repair DNA to ensure their survival [62]. FMM determines the net amount of DNA damage, detected as single- and double-stranded DNA breaks and alkali labile sites, which result from both concomitant damage and repair [46,48]. This is a good indicator of environmental stress in marine invertebrates [46,48]. If *C. hyperboreus* cannot survive in the Skagerrak and is a non-native species transported by ocean currents, as suggested [24,27,32], one would expect the species to have high levels of DNA damage due to stress. Conversely, *C. hyperboreus* exhibits overall less DNA damage than *C. finmarchicus* and *M. longa*, suggesting that its biology is adapted to life in the Oslofjord. Temperature can increase the rate of DNA damage either by increasing ROS production through metabolism or by decreasing DNA repair by altering enzyme activity [37,63]. Here, we indeed find that DNA damage increases with temperature in all three species without interspecific differences.

Our results suggest the presence of an isolated population of *C. hyperboreus* in the Skagerrak that is adapted to warmer temperatures than Arctic conspecifics. The literature on *C. hyperboreus* outside the Arctic is scarce but supports our hypothesis. First, *C. hyperboreus* from the Oslofjord have smaller genomes than in the Arctic, which is generally associated with adaptation to warmer temperatures [25]. Secondly, there is a discontinuity in the range of the species. Indeed, *C. hyperboreus* are negligible in the Norwegian and North Seas. Yet, abundances in the Skagerrak, off Kristiansand, can be as high as 3.2 g DW m^-2^ [28,29]. These abundances are similar to those in much of the Arctic Ocean, such as the Amundsen, Canada, Makarov, and Nansen basins, as well as Svalbard waters, which are typically 2–3 g DW m^-2^ [33–36]. It is unlikely that *C. hyperboreus* would concentrate in Skagerrak only through advection via oceanic currents as it is often suggested [24,27,32]. All surface currents outflow from Skagerrak, the only current coming from the North is the Norwegian trench inflow in the deep (< 200m) [64]. The Norwegian trench inflow transports overwintering *C. finmarchicus* from the Norwegian sea into Skagerrak during the winter [65,66]. However, *C. hyperboreus* are considered unable to overwinter at such southern latitudes [24,27,32].

*C. hyperboreus* is a key species in Boreal and Arctic ecosystems as the main prey of many fish species and seabirds, and by storing carbon in the deep sea via overwintering migration [15,17,18]. Models predict that the species’ range will shift northward with OW, losing up to 70 km per decade in the coming years [14], with large consequences for ecosystems. Here we show that local adaptations to warmer waters likely exist in the Skagerrak and should be considered when projecting the future of *C. hyperboreus*. Further studies directly comparing the ecology, genetics, connectivity and stress tolerance threshold of *C. hyperboreus* between Skagerrak and Arctic populations would greatly improve our knowledge of local adaptations and global change. This is crucial since the existence of local adaptations can constitute a reservoir of resistant genotypes which can benefit the persistence of species in the face of environmental changes [38,39].

## Authors’ contributions

M.L.: conceptualization, data curation, formal analysis, investigation, methodology, validation, writing - original draft; A.E.T.S.: investigation, methodology, validation; N.T.: investigation, methodology, validation; H.R.: investigation, methodology, validation; K.V.D.: conceptualization, investigation, methodology, validation, supervision, funding acquisition, project administration, writing - review and editing.

All authors gave final approval for publication and agreed to be held accountable for the work performed therein.

## Acknowledgments

The authors thank research groups at the University of Oslo and the Scottish Association for Marine Sciences for assistance with field sampling and DNA damage analyses.

