## Supplementary Information for "The Arctic copepod *Calanus hyperboreus* is more tolerant to marine heatwaves than temperate copepods in the Oslofjord"

### Tables

**Supplementary Table 1.** Number of copepods analysed for DNA damages for each species and each temperature.

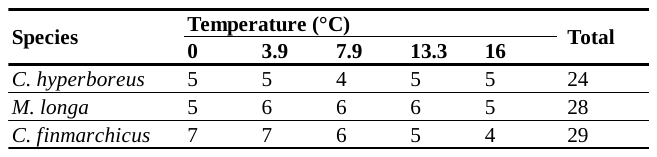

**Supplementary Table 2.** Results of the Cox proportional-hazards model testing the survival of three species of copepods from the Oslofjord as a function of temperature. The global model tests the effects of species, temperature and their interaction but is unbalanced (n = 245 for *Metridia longa*, n = 105 for *Calanus finmarchicus* and n = 35 for *Calanus hyperboreus*). The individual models test the effect of temperature for each species alone, with balanced data.

| **Model** | **Factor** | **Degree of freedom** | **Chi-squared statistic** | **p value** |
| --- | --- | --- | --- | --- |
| Global | Temperature | 1 | 377.7 | < 2.200 10^-16^ |
|  | Species | 2 | 20.7 | 3.236 10^-5^ |
|  | Temperature*Species | 2 | 62.2 | 3.133 10^-14^ |
| *M. longa* | Temperature | 1 | 247.8 | < 2.200 10^-16^ |
| *C. finmarchicus* | Temperature | 1 | 120.8 | < 2.200 10^-16^ |
| *C. hyperboreus* | Temperature | 1 | 37.3 | 1.017 10^-9^ |

**Supplementary Table 3.** Results of the type II ANOVA testing the DNA damages of three species of copepods from the Oslofjord as a function of temperature. The effects of species, temperature and their interaction are tested

| **Factor** | **Degree of freedom** | **F statistic** | **p value** |
| --- | --- | --- | --- |
| Temperature | 1 | 4.4827 | 0.03774 |
| Species | 2 | 32.3158 | 1.049 10^-10^ |
| Temperature*Species | 2 | 0.3340 | 0.71721 |
| Residuals | 71 |  |  |

**Supplementary Note 1.**

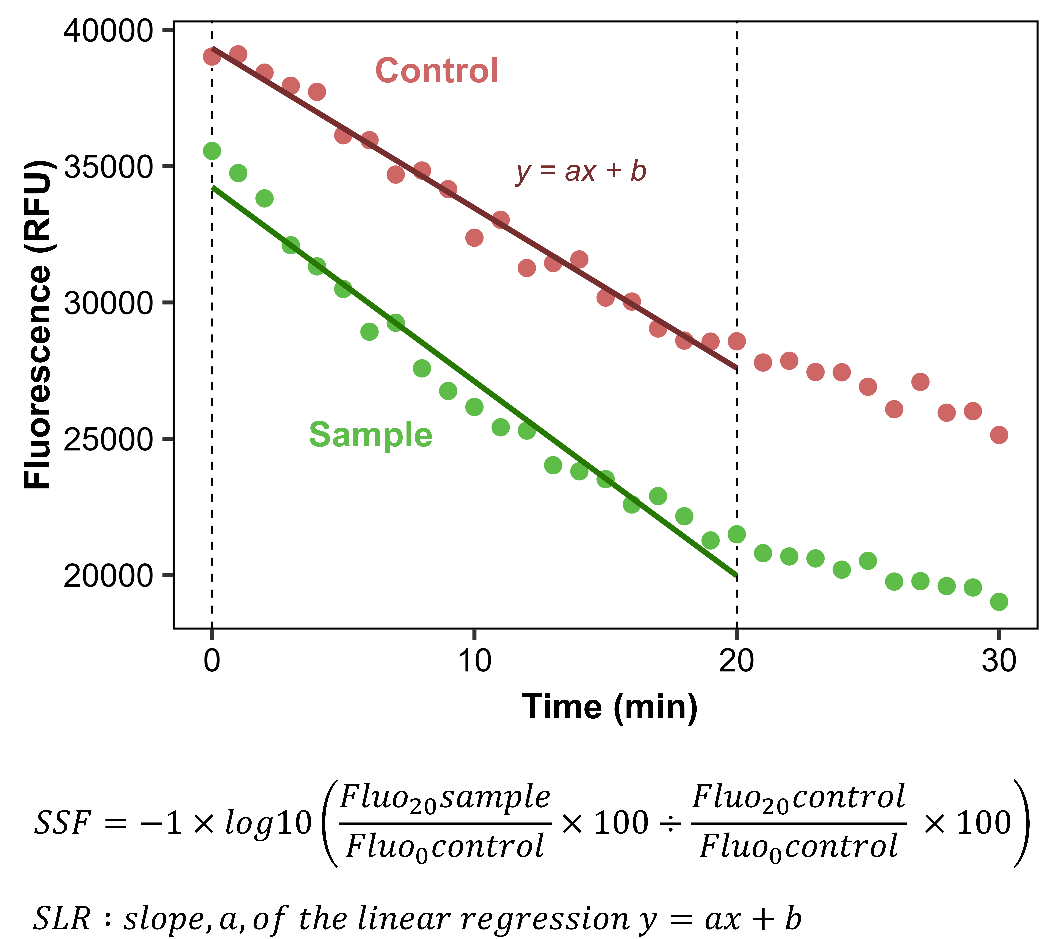
The formulas used to calculate the *Strand Scission Factor* (SSF) and the *Slope of the Linear Regression* (SLR) are shown in the figure below. *Fluo_0_control*, *Fluo_20_control*, *Fluo_0_sample*, *Fluo_20_sample* are the fluorescence values at 0 and 20 minutes in the control and the sample respectively. To determine the SLR, a linear regression model is fitted between 0 and 20 minutes. We only kept the linear models that were statistically significant and met the assumptions of normality and homoscedasticity of the residuals. The SSF has the disadvantage of relying on controls, i.e. 1 or 2 copepods, that are highly variable between microplates. In addition, the calculation of the SSF is based on only two fluorescence points, which induces some variability if there are outliers in the fluorescence measurements at 0 and 20 minutes, due to microplate reader imprecisions. The SLR is independent from control and relies on 21 fluorescence values for its calculation. The example below comes from real data of DNA damages measured on *Calanus hyperboreus* samples at 8 °C (control) and 12 °C (sample). In our data, SLR is correlated with SSF (R^2^ = 0.54, Figure S2) and thus an increase in SLR shows an increase in DNA damages. This is illustrated in the figure below with all SLR calculated on *M. longa*, *C. finmarchicus* and *C. hyperboreus* for all temperatures, modeled as a function of SSF with a linear regression.

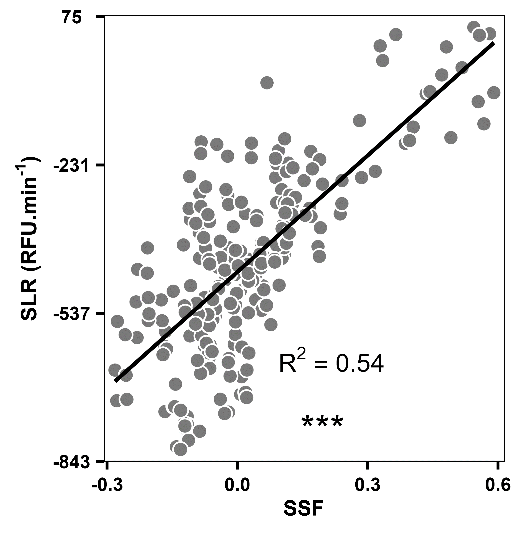
